# Multimodal Predictive Modeling: Scalable Imaging Informed Approaches to Predict Future Brain Health

**DOI:** 10.1101/2024.05.29.596506

**Authors:** Meenu Ajith, Jeffrey S. Spence, Sandra B. Chapman, Vince D. Calhoun

**Author notes:** Correponding author, *Email address:* (Meenu Ajith).

## Abstract

**Background:** Predicting future brain health is a complex endeavor that often requires integrating diverse data sources. The neural patterns and interactions identified through neuroimaging serve as the fundamental basis and early indicators that precede the manifestation of observable behaviors or psychological states.

**New Method:** In this work, we introduce a multimodal predictive modeling approach that leverages an imaging-informed methodology to gain insights into future behavioral outcomes. We employed three methodologies for evaluation: an assessment-only approach using support vector regression (SVR), a neuroimaging-only approach using random forest (RF), and an image-assisted method integrating the static functional network connectivity (sFNC) matrix from resting-state functional magnetic resonance imaging (rs-fMRI) alongside assessments. The image-assisted approach utilized a partially conditional variational autoencoder (PCVAE) to predict brain health constructs in future visits from the behavioral data alone.

**Results:** Our performance evaluation indicates that the image-assisted method ex-cels in handling conditional information to predict brain health constructs in subsequent visits and their longitudinal changes. These results suggest that during the training stage, the PCVAE model effectively captures relevant in-formation from neuroimaging data, thereby potentially improving accuracy in making future predictions using only assessment data.

**Comparison with Existing Methods:** The proposed image-assisted method outperforms traditional assessment-only and neuroimaging-only approaches by effectively integrating neuroimaging data with assessment factors,

**Conclusion:** This study underscores the potential of neuroimaging-informed predictive modeling to advance our comprehension of the complex relationships between cognitive performance and neural connectivity.

**Highlights:**

- Multifaceted perspective for studying longitudinal brain health changes.
- Showcases the versatility of methodologies through assessment-only, neuroimaging-only, and image-assisted predictive approaches.
- Provides predictive insights by revealing the neural patterns corresponding to alterations in behavior.

**Graphical Abstract:** 

## 1. Introduction

Brain health is conceptualized as the primary domain of health, with mental health and psychosocial well-being falling under its umbrella as sub-ordinate components Chapman et al. (2022). According to the World Health Organization (WHO), brain health involves the continuous promotion of optimal brain development and fitness throughout one’s lifespan, incorporating cognitive health, emotional well-being, and the importance of social connections and purpose Organization et al. (2022). Therefore, it is imperative to devise strategies to enhance and optimize brain health, thereby enhancing the quality of life for the population and mitigating the burden of neurological diseases and mortality Hachinski et al. (2021).

The multifaceted nature of brain health emphasizes the need for a more holistic approach, departing from current fragmented assessments. This holistic perspective involves a dynamic interplay of cognitive, emotional, social, and lifestyle factors, with brain health as a crucial determinant Avan et al. (2022). Alterations in neural structure and activity serve as pivotal regulators of behavior, influencing individuals’ responses to stimuli and their navigation of daily life. In a comprehensive analysis Lee et al. (2024), 478 distinct methods of brain health measurement were identified, and among them, 268 (56.1%) were utilized only once. The remaining 210 methods incorporated measurements from various sources, including imaging, biological, clinical, mental health, and cognitive tests. Among these approaches, those based on imaging were the most prevalent for assessing brain health. The majority of unimodal MRI-based imaging techniques for assessing brain health relied on estimating volumes of grey and white matter in specific regions, notably the hippocampus and the entire brain, along with identifying white matter hyperintensities Hu et al. (2021) and fractional anisotropy. Additionally, the Trail Making Test (TMT) Guo (2022) and the Mini-Mental Status Examination (MMSE) Arevalo-Rodriguez et al. (2015) emerged as the two most commonly used cognitive assessment methods. These cognitive tests and neuropsychological assessments play a crucial role in predicting brain health outcomes, particularly in conditions characterized by cognitive impairment Battista et al. (2017). These assessments encompass tasks related to memory, attention, and executive function. Several cognitive assessments have been modified for utilization within task-based fMRI scans, enhancing the feasibility of integrating imaging with cognitive evaluation in assessing brain health Guo (2022). To enhance predictive capabilities, machine learning and statistical modeling techniques are frequently employed, utilizing unimodal data sources Nemesure et al. (2021). These models can identify patterns and relationships that contribute to predicting brain health outcomes.

Relying exclusively on cognitive testing for brain health assessment carries costs and sensitivity limitations. Moreover, there is a risk of bias impacting prediction reliability, and these assessments may overlook crucial neurobiological factors affecting brain health Lira et al. (2022). On the contrary, the primary use of MRI in assessing brain health is constrained by its significantly high cost. Moreover, depending solely on data from a single modality can result in a limited perspective of the complex nature of brain. In predictive modeling, integrating multimodal data is a powerful tool for understanding complex systems. This study utilizes innovative strategies and diverse data sources, particularly static functional network connectivity (sFNC) data, to uniquely integrate neuroimaging insights, revealing subtle links between neural activity and shifts in brain health over time. This novel method minimizes reliance on neuroimaging data for prediction by learning concealed hidden imaging-behavior relationships during training, thus eliminating the need for imaging data in predicting new data. This has the advantage of reducing the necessity for continuously gathering extensive imaging data, which can be resource-intensive and time-consuming. By decreasing dependence on neuroimaging data for prediction, the model becomes more applicable for real-world use. It broadens the scope of advanced predictive modeling across various fields such as healthcare, psychology, and neuroscience, where access to neuroimaging data may be limited. Additionally, it enhances efficiency by eliminating the need for continual training on extensive datasets, which is particularly crucial in time-sensitive scenarios like clinical decision-making or cognitive assessment.

This predictive framework relies on three distinct methodologies. Firstly, an assessment-only approach that employs support vector regression (SVR). Secondly, a neuroimaging-only approach which utilizes random forest (RF). Finally, the proposed image-assisted method involves a partially conditional variational autoencoder (PCVAE). We conduct two types of prediction scenarios: one to forecast future brain health and the other to predict changes in brain health over time. The image-assisted method integrates the sFNC matrix from neuroimaging data with assessment factors of brain health. This approach excels in handling conditional information to predict brain health constructs in subsequent visits and their longitudinal changes. Understanding the correlation between functional network connectivity within the brain and specific assessment measures of brain health may enable early identification or anticipation of neurological conditions, as well as fostering enhanced cognitive function, mental resilience, and overall well-being through targeted interventions. The proposed image-assisted method establishes a fundamental framework for assessing brain health and capturing the complex connections between brain networks, as well as their associations with various psychosocial factors. The main contributions of this paper are as follows:

- Multifaceted Predictive Framework Integration: The paper introduces a comprehensive predictive framework that intricately combines assessment data with insights from neuroimaging. This integration offers a deep understanding of the relationship between latent brain health constructs and neural patterns, providing a holistic perspective for studying longitudinal brain health changes.
- Practical Alternative to Costly Neuroimaging: By eliminating the need for neuroimaging data during testing, the model becomes more resource-efficient, requiring fewer computational resources and time for inference. This ensures that the model remains applicable in real-world scenarios where access to neuroimaging data may be challenging due to factors like cost, accessibility, or ethical concerns.
- Neural-Behavioral Correlations: Through the image-assisted approach, the study delves into the complex interplay between neural activity and behavioral changes over time. By utilizing neuroimaging data alongside assessment factors, it aims to unravel correlations, potentially uncovering how neural patterns correspond to improvement or losses in behavior and paving the way for predictive insights into these changes.
- Methodological Framework Advancements: The paper introduces and describes the application of PCVAE for the prediction of behavioral constructs. This novel framework allows for the effective integration of neuroimaging data and assessment factors. It also facilitates learning joint representations and ultimately enables improved predictive values of behavioral change trajectories across multiple visits.

## 2. Materials and Methods

### 2.1. UTD Data Acquisition and Preprocessing

The assessment data used for testing was obtained from the BrainHeath Project led by the Center for BrainHealth at the University of Texas at Dallas (UTD) Chapman et al. (2021). Participants were recruited through word-of-mouth and email advertisements, focusing on generally healthy adults. To determine eligibility, potential participants completed an online screening form. The inclusion criteria required participants to be 18 years of age or older, have internet access, and be proficient in English. The exclusion criteria were being under 18, having a diagnosed neurological or psychotic disorder, uncontrolled psychiatric disorder, history of brain injury, or any uncontrolled health issues. Moreover, participants were not excluded based on general health risk factors such as obesity, diabetes, or autoimmune conditions. In our study, we analyzed data from a total of 158 participants who initially completed baseline assessments for the Visit 1 Brain Health Index (BHI) and subsequently completed the Visit 2 BHI after 3 months of engaging in online training and coaching interventions.

Initially, participants underwent a series of online assessments targeting four essential domains of brain health: (1) cognition, (2) well-being, (3) social interaction, and (4) daily life. The cognitive assessments cover a range of measures that pertain to the comprehension of complex texts, ensuring that any limitations that may restrict accurate evaluation are avoided. The cognitive evaluation focuses on diverse aspects of advanced cognitive abilities, including reasoning, innovation, strategy, and memory. The well-being evaluation delves into an individual’s emotional well-being, encompassing various aspects such as quality of life, level of happiness, stress levels, sadness, and emotional resilience. The social interaction evaluation examines an individual’s level of social engagement and the quality of their relationships, specifically exploring their perceptions of social support networks and the meaningfulness of their social interactions. Lastly, the daily life evaluation assesses the intricacy and breadth of an individual’s daily responsibilities, habits, and challenges, aiming to understand how individuals optimize their life circumstances and routines. Table. 1 presents the assessment measures and their corresponding categories in the BrainHeath Project dataset.

**Table 1:** Assessment measures of UTD dataset for BHI.

| Brain systems | Assessments | Measurement scales/task |
| --- | --- | --- |
| Cognition | Strategic attention | Visual selective learning task Hanten et al. (2007) |
|  | Innovation | Pictures |
|  | Innovation | Test of strategic learning (High level interpretation) Vas et al. (2015) |
|  | Abstraction | Proverbs |
|  | Integrated reasoning | Test of strategic learning (High-level summary of text) Vas et al. (2015) |
|  | Memory of detail | Test of strategic learning Vas et al. (2015) |
| Well-being | Happiness | Oxford happiness questionnaire Hills (2002) |
|  | Resilience | Connor-Davidson resilience scale Schwarzer and Jerusalem (1995) |
|  | Depression | Depression, anxiety, stress scale Lovibond and Lovibond (1995) |
|  | Anxiety | Depression, anxiety, stress scale Lovibond and Lovibond (1995) |
|  | Stress | Depression, anxiety, stress scale Lovibond and Lovibond (1995) |
|  | Life satisfaction | Quality of life scale Burckhardt et al. (2003) |
| Interaction | Social support | Social support survey index Sherbourne and Stewart (1991) |
|  | Social engagement | Social brain health scale Allen et al. (2020) |
|  | Compassion | Adapted from Strauss et al. (2016); Johnson (2018) |
| Daily life | Sleep | Pittsburgh sleep quality index Buysse et al. (1989) |
|  | Metabolic equivalents | Cardiorespiratory fitness estimate questionnaire Jurca et al. (2005) |
|  | Outlook | Brain health appraisal questionnaire |
|  | Meaningful activities | Engagement in meaningful activities survey Eakman (2011) |

In prior research Chapman et al. (2021), we utilized an exploratory factor model to understand the relationships among different assessments. This model aimed to capture the shared variance or common factors among these measures, allowing us to understand which factors contribute most significantly to changes in brain health over time. The factor analysis resulted in a 3-factor solution, and the model’s output provided appropriate variable weights that could transform individual assessments into these identified factors. Hence, normalized factor loadings were acquired for all 20 assessment measures across the three identified factors. We multiplied the assessment measures by their respective factor loadings to compute factor scores, such as Factor 1, Factor 2, and Factor 3, for each subject. Here Factor 1 is labeled as connectedness to people and purpose due to its encompassment of metrics associated with social engagement and behavioral aspects such as mental resilience and life purpose/satisfaction. Factor 2, termed emotional balance, primarily encompasses mood evaluations within the domain of well-being, focusing notably on depression, anxiety, stress, and happiness. On the other hand, Factor 3, referred to as clarity, is primarily characterized by assessing strategic attention, innovation, integrated reasoning, and memory for complex information. The BHI was calculated by aggregating these factors to generate an overall index score reflecting a composite brain health score. In this work, we aim to predict the three identified factors at time point 2 and their changes over 3 months, which are pivotal in assessing individuals’ brain health status.

For the 4-D preprocessed rs-fMRI data, we applied a fully automated spatially constrained Independent Component Analysis (ICA) process known as NeuroMark Du et al. (2020). We employed a template consisting of reproducible independent components (ICs), which were created through the spatial alignment of correlated group-level ICs from two extensive healthy control fMRI datasets with large sample sizes (N *>* 800): the Genomics Superstructure Project (GSP) and the Human Connectome Project (HCP). The Neuromark fMRI 1.0 template was individually applied to each subject and served as priors for the spatially constrained ICA algorithm. This process aimed to identify 53 functionally relevant resting-state networks (RSNs) for each individual, emphasizing maximum spatial independence. Each RSN comprises a spatial map and its corresponding time course (TC). The sFNC reflecting interactions between any two networks was ultimately derived by computing Pearson correlations between the time courses. The RSNs were categorized into seven domains based on functional similarity, namely sub-cortical (SC), auditory (AUD), sensory-motor (SM), visual (VIS), cognitive control (CC), default mode (DM), and cerebellar (CB).

### 2.2. Prediction Framework

This approach revolves around identifying hidden variables (latent constructs) from the observed assessment data. Three such latent variables or factors have been calculated and are the focal points of this study. The methodology involves using these identified factors from an initial visit to predict or estimate the same latent constructs in a subsequent visit, indicating a longitudinal investigation of changes across time. Additionally, the study incorporates neuroimaging data to delve into the relationship between neural activity and these hidden behavioral constructs, aiming to uncover connections between brain processes and changes in behavior over time. This multifaceted approach aims to unravel how neural patterns correspond to alterations in behavioral tendencies. It seeks to shed light on the interplay between brain activity and behavioral changes across different time points, potentially predicting these changes.

This research methodology is based on predicting three derived factors and their temporal changes over 3 months, utilizing data gathered at baseline (Visit 1) and after intervention (Visit 2) through an image-assisted approach. The image-assisted approach is contrasted with assessment-only and neuroimaging-only methods to conduct additional performance evaluation. All three approaches analyze two scenarios: one to forecast Visit 2 factor scores and the other to predict the change in factors (Visit 2-Visit 1). The image-assisted method incorporates both sFNC data and factors, utilizing a specialized version of variational autoencoders, a model tailored for multimodal data, to predict the factors from Visit 2 and their changes. Consequently, this study aims to assess the predictive capability of multimodal data and explore their integration with neuroimaging data in forecasting brain health constructs across multiple visits.

A PCVAE is employed in the image-assisted method primarily due to its capability to handle conditional information effectively. In this approach to forecasting the Visit 2 factors, the sFNC features are utilized as inputs for the encoder part, with the factor scores serving as conditioning variables. The PCVAE allows the incorporation of this additional information into the generation process of latent representations. The encoder compresses these features into a latent vector that encapsulates both the sFNC and factor score information. Subsequently, the decoder reconstructs this combined representation to predict the factor scores. Essentially, the PCVAE framework facilitates learning a joint representation of sFNC features and factor scores. This enables the model to effectively leverage the relationship between these features for accurate predictions. During training, both sFNC features and factor scores are fed into the model, but during testing, only the factor scores are used. The encoder consists of densely connected layers with 16 and 4 nodes, generating a latent vector of dimension 2. The decoder mirrors this architecture with layers of 16 and 4 nodes, culminating in a final dense layer responsible for predicting the factor scores. This process enables the model to capture and reconstruct relationships between sFNC features and factor scores, ultimately predicting factor scores accurately. Additionally, the PCVAE model used leaky rectified linear unit (Leaky ReLU) Maas et al. (2013) non-linearity in its convolutional layers. It is characterized by allowing a small, non-zero gradient for negative inputs, unlike the traditional ReLU function, which zeros out negative values. In Leaky ReLU the alpha parameter determines the slope of the function for negative inputs. In this case, the optimum alpha value was found to be 0.1 after hyperparameter tuning. Hence, instead of setting negative values to zero, they are multiplied by 0.1, allowing for a slight gradient, which can prevent the issues of dying neurons often associated with ReLU activations. The architecture of the proposed model is shown in Fig. 1. The same model is employed to predict the change in factors, incorporating the difference in sFNC (Visit 2-Visit 1) alongside the Visit 2 factors.

**Figure 1:**
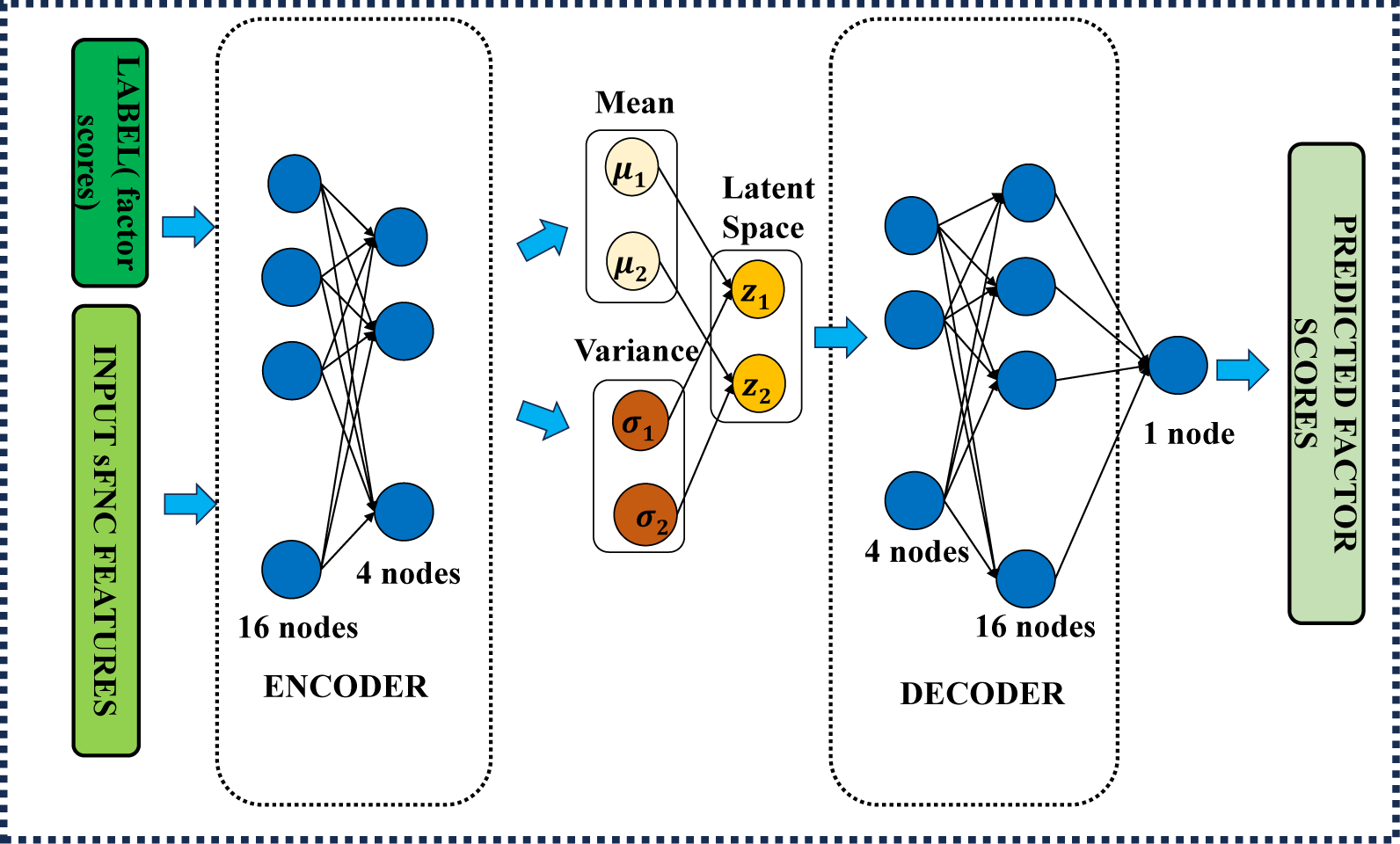
Architecture of the proposed PCVAE for the image-assisted model.

Furthermore, optimizing the network’s parameters during training involved using the Adam optimizer Kingma and Ba (2014). Specifically, this optimizer was employed for 1500 iterations, utilizing a learning rate of 0.001 and a smaller batch size of two. A reduced batch size facilitates more frequent parameter updates throughout the training, potentially enabling finer adjustments in the model. For the training approach, a ten-fold cross-validation method was applied to the dataset. This method partitions the dataset into ten distinct folds. During the training phase, nine of these folds were iteratively utilized to train the model, while the remaining fold served as the test set. This process was repeated ten times, rotating through each fold as the test set, allowing for a comprehensive evaluation of the model’s performance across various subsets of the data.

Meanwhile, the assessment-only approach exclusively depends on factors for prediction and employs an SVR model. This model utilizes a radial basis function (RBF) kernel with specific hyperparameters set as follows: C = 1.0 and epsilon = 0.1. Therefore, by using an RBF kernel, the SVR model can better handle complex data structures and capture the nonlinear relationship between the variables. The C parameter, set at 1.0, determines the trade-off between minimizing training error and maximizing the decision boundary. At the same time, epsilon, fixed at 0.1, establishes the tolerance margin for errors within the model. In contrast, the neuroimaging-only approach relies solely on sFNC data. It utilizes an RF regression model, where the number of estimators is fixed at 10, determining the quantity of decision trees constructed during the training phase.

## 3. Results

The neuroimaging features of each participant are represented by 53 × 53 sFNC matrices, showcasing the connectivity strengths among different RSNs. As these matrices are symmetrical, we extract only the upper triangular section and flatten it. This process yields the final flattened feature vector for the sFNC data, sizing up to 1378 × 1. For all models, standardization is done as a preprocessing step to transform features to have a mean of 0 and a standard deviation of 1. This process helps bring different features to a similar scale, preventing certain features from dominating the learning process due to their larger magnitudes. Following this, a feature selection was made by calculating the correlation coefficients between the flattened sFNC values from Visit 1 and the factors from Visit 1. Calculating correlation coefficients between features helps identify relationships between these variables. Here, we identify the top correlated features based on a threshold and select those features from the flattened sFNC values. Setting a threshold for correlation coefficients helps identify which features are more strongly associated with the factors. This step helps in reducing dimensionality, computational complexity, and potentially overfitting by including only the most informative features for modeling. Finally, these features are given as input to the PC-VAE model. We employ connectograms to visually represent the strongest correlations between the flattened sFNC and the factors. Connectograms are powerful tools used to illustrate and analyze the connections between different regions of the brain. These visual representations offer a comprehensive view of the intricate network of neural pathways and connections within the brain. Here, the connectogram analysis is conducted using the ICs derived from the spatially constrained ICA of the rs-fMRI data.

The given connectogram analysis, depicted in Fig. 2 (a), (b), and (c), serves as a pivotal method to visually showcase the neurological differences linked to the three factors. The connectograms offer a comprehensive portrayal of neural pathways, aiding in understanding and highlighting distinctive neural connectivity patterns associated with each factor. Within this visualization, the blue lines between the ICs indicate a negative correlation, while the yellow lines denote strong positive connections. The transparency of the lines reflects the strength of these connections. Positive correlations among the ICs imply that when one brain area is active, the other tends to be active as well, indicating functional coherence or synchronization between these regions. Conversely, a negative correlation indicates that when one brain area is active, the other tends to be less active or inactive, indicating functional antagonism or suppression between these regions.

**Figure 2:**
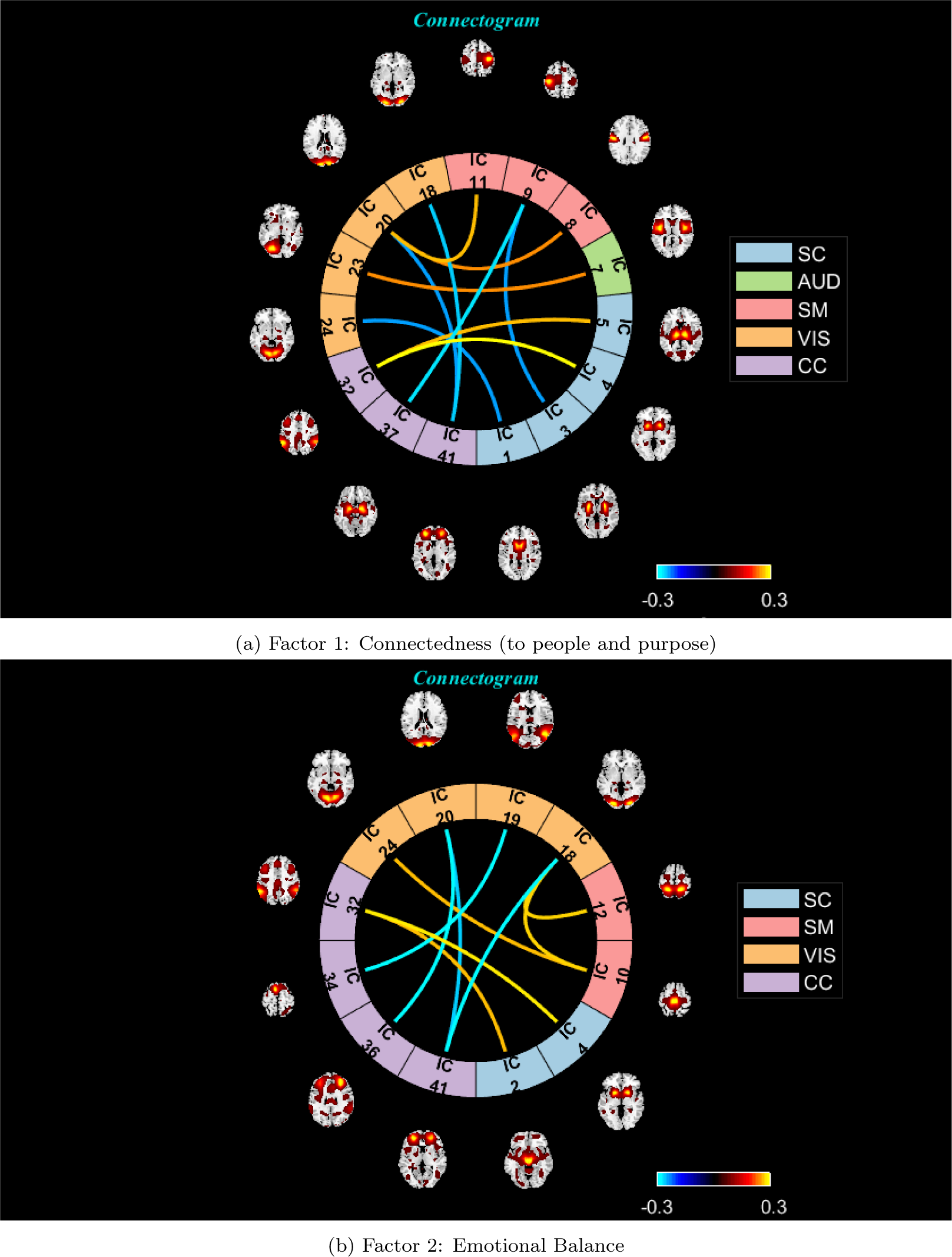
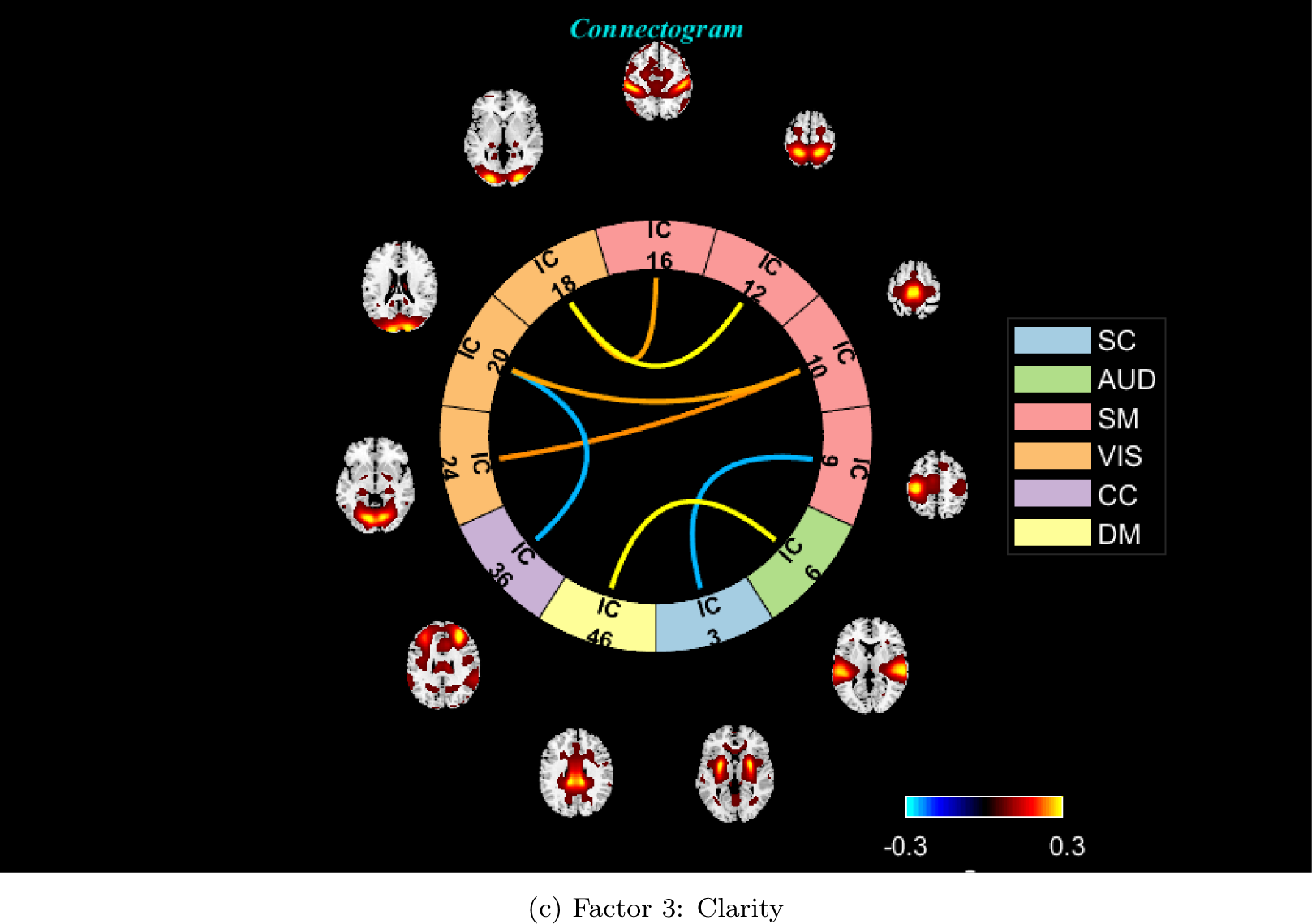
The connectogram visually represents the connections between different resting-state networks (RSNs) based on their functional correlations. This depiction illustrates the relationship between the factors and the sFNC matrix. Yellow hues connecting the independent components (ICs) highlight brain areas exhibiting positive correlations among features, while blue lines connecting the ICs indicate regions displaying negative correlations among these features. ICs play a crucial role in identifying and characterizing RSNs. The RSNs were categorized into seven domains: subcortical (SC), auditory (AUD), sensory-motor (SM), visual (VIS), cognitive control (CC), default mode (DM), and cerebellar (CB).

The connectogram for factor 1 exhibited significantly positive values in the cognitive control-subcortical (CC-SC) domains. This indicates strong connectivity between brain areas responsible for decision-making, attention, etc., and subcortical regions such as the thalamus and the basal ganglia. On the other hand, negative values are observed in connection pairs such as the cognitive control-sensorimotor (CC-SM) and cognitive control-visual (CC-VIS) domains. Negative values signify weaker associations between brain regions responsible for cognitive control and those involved in sensorimotor or visual processing. Furthermore, the connectogram pattern reveals an overlap between factor 2 and factor 1 in terms of positive correlations within the cognitive control-subcortical (CC-SC) domains. Stronger correlations between cognitive control regions and subcortical brain areas imply decision-making efficiency, emotional regulation, and more effective responses in complex situations by integrating information from subconscious brain regions. Additionally, factor 2 demonstrates enhanced connectivity between visual processing and sensorimotor functions. This indicates potential improvements in co-ordinating visual perception with movement-related activities. In contrast, there is reduced connectivity between cognitive control and visual processing regions.

Finally, the connectogram depicting factor 3 displayed notably strong connections in the visual-sensorimotor (VIS-SM) and default mode-auditory (DM-AUD) domains. This heightened connectivity between the default mode network and auditory regions denotes a potentially stronger association between internal thoughts, self-referential thinking, and auditory perception. Additionally, comparable to factors 1 and 2, factor 3 demonstrated decreased correlations within the cognitive control-visual (CC-VIS) domain.

### 3.1. Evaluation of Prediction Errors for Post-Intervention Factors and Changes

Table. 2 assesses the post-intervention prediction performance, specifically predicting the factors for Visit 2 using the provided test data. The evaluation involves three models: SVM, RF, and PCVAE. Each of these models is evaluated under three distinct scenarios for factors labeled 1 through 3. Initially, predictions are based solely on Visit 1 data for the respective factor. Subsequently, a model is employed utilizing only sFNC data from Visit 1. Finally, predictions are refined by integrating sFNC data from Visit 1 with factor-related data. The performance of each model is evaluated using metrics such as mean squared error (MSE), mean absolute error (MAE), and the coefficient of determination (R-squared or R2) metrics. Lower values of MSE and MAE indicate better performance, while a higher R2 value suggests a better fit of the model to the data. A global Min-Max scaling was applied to the sFNC and factor scores to ensure that the values were within the same range. Subsequently, both the predicted and actual values underwent an inverse transformation to restore them to their original range between 300 and 700. The R2 values remained consistent regardless of whether the predicted and actual values were utilized before or after the inverse transformation. However, the values presented in Table. 2 and 3 were computed before the inverse transform, resulting in relatively smaller values.

**Table 2:** Performance evaluation of the post-intervention (Visit 2) factor prediction using the proposed approaches.

| Factors | Prediction | Train data | Test data | Model | MSE | MAE |
| --- | --- | --- | --- | --- | --- | --- |
| Factor 1 | Factor 1 Visit 2 | Factor 1 Visit 1 | Factor 1 Visit 1 | SVM | 0.62 | 0.61 |
|  |  | sFNC Visit 1 | sFNC Visit 1 | RF | 0.85 | 0.70 |
|  |  | sFNC Visit 1 +<br>Factor 1 Visit 1 | Factor 1 Visit 1 | PCVAE | 0.02 | 0.13 |
| Factor 2 | Factor 2 Visit 2 | Factor 2 Visit 1 | Factor 2 Visit 1 | SVM | 0.90 | 0.71 |
|  |  | sFNC Visit 1 | sFNC Visit 1 | RF | 0.86 | 0.77 |
|  |  | sFNC Visit 1 +<br>Factor 2 Visit 1 | Factor 2 Visit 1 | PCVAE | 0.73 | 0.67 |
| Factor 3 | Factor 3 Visit 2 | Factor 3 Visit 1 | Factor 3 Visit 1 | SVM | 0.89 | 0.74 |
|  |  | sFNC Visit 1 | sFNC Visit 1 | RF | 0.64 | 0.67 |
|  |  | sFNC Visit 1 +<br>Factor 3 Visit 1 | Factor 3 Visit 1 | PCVAE | 0.03 | 0.14 |

**Table 3:**
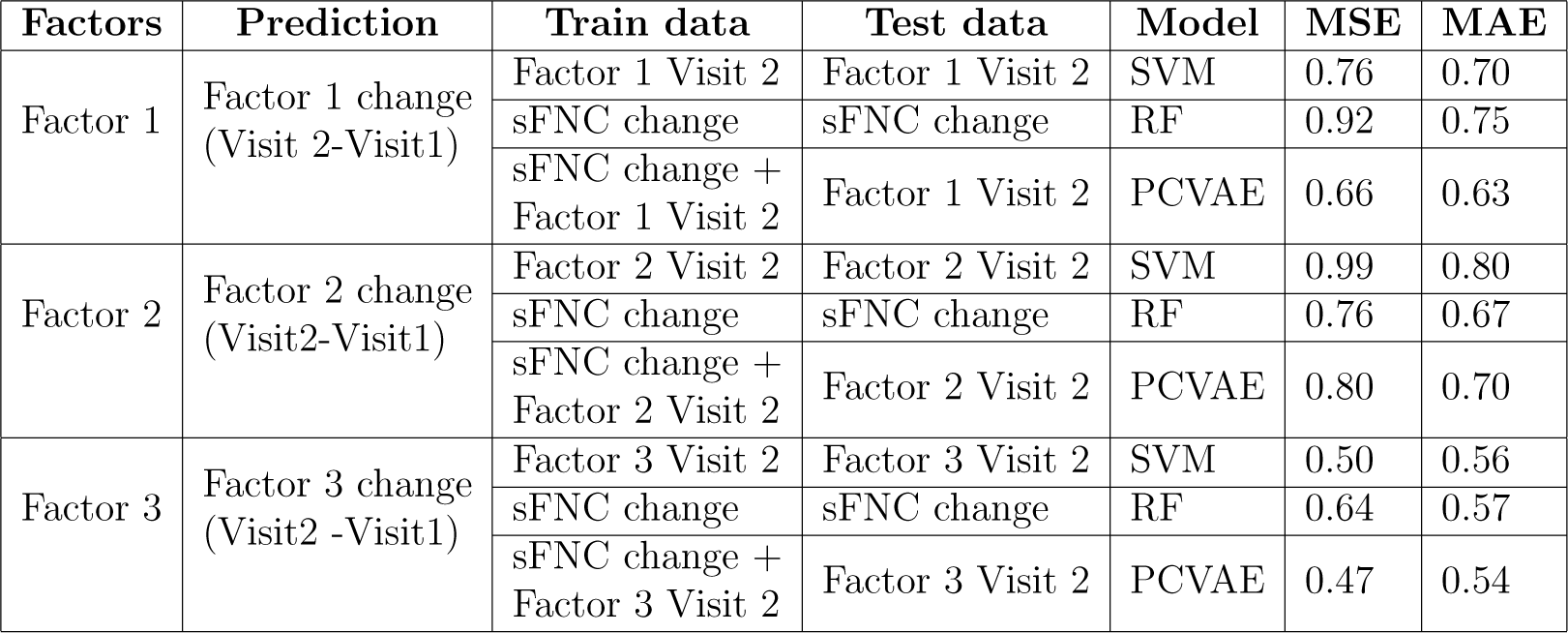
Performance evaluation of the change in (Visit 2-Visit 1) factors using the proposed approaches.

For the prediction of all three factors, PCVAE significantly outperforms SVM and RF in terms of both MSE and MAE. The assessment-only model demonstrates the second-best performance for Factor 1, whereas the neuroimaging-only model exhibits the second-best performance for Factors 2 and 3. The SVR model shows a moderate predictive error, with MSE ranging from 0.62 to 0.90 and MAE ranging from 0.61 to 0.74. Similarly, the RF model exhibits an MSE ranging from 0.64 to 0.86 and an MAE ranging from 0.67 to 0.77. In contrast, the image-assisted model provides the most accurate predictions, with MSE ranging from 0.02 to 0.73 and MAE ranging from 0.13 to 0.67. Notably, the PCVAE model demonstrates a substantial improvement in predictive accuracy with the inclusion of sFNC Visit 1 data alongside factor-related information. This highlights the efficacy of incorporating neuroimaging data to enhance the precision of post-intervention predictions using the PCVAE model.

Table. 3 presents the performance evaluation of different approaches for predicting the change in factors between Visit 2 and Visit 1. Analyzing the results, we observe variations in the performance of the prediction models across different factors. For Factor 1, the PCVAE approach outperforms SVM and RF in terms of both MSE and MAE. Similarly, for Factor 2, while SVM performs relatively poorly compared to RF and PCVAE, the latter two models demonstrate closer performance. For Factor 3, all models show relatively similar performance, with PCVAE having the lowest MSE and MAE. Overall, the PCVAE approach consistently shows competitive performance across all factors, indicating its effectiveness in predicting the changes between Visit 2 and Visit 1.

Fig. 3 displays the regression plots generated by the image-assisted model, illustrating its predictions for Visit 2 factors and the change in factors (Visit 2 -Visit 1) across factors 1, 2, and 3. The left pane, depicted in panels (a), (c), and (e) of Fig. 3, employs the PCVAE model to forecast Visit 2 factor scores by combining both sFNC data from Visit 1 and factors from Visit 1. Notably, Factor 1 demonstrates the highest R2 score compared to Factors 2 and 3. The actual scores for Factor 1 spanned between 300 and 750, while the predicted scores ranged from 350 to 700, suggesting that the model’s predictions fall within a relatively close range of the actual scores. Likewise, the right panel illustrated in Fig. 3 (b), (d), and (f) utilizes the PCVAE model, incorporating changes in sFNC data and factors from Visit 2 to forecast changes in factors across visits. The change in Factor 3 exhibits the highest R2 score, with the regression lines appearing tightly clustered and closely aligned to the ideal predictive trajectory. Consequently, this indicates a stronger and more consistent relationship between the actual and predicted scores compared to Factors 1 and 2. Hence, the image-assisted model offers insights into how factors evolve between visits, providing valuable information for understanding longitudinal trends and potentially uncovering underlying patterns of change. Moreover, the incorporation of image data alongside the factors showcases the potential benefits of utilizing a broader range of information sources in predictive modeling scenarios.

**Figure 3:**
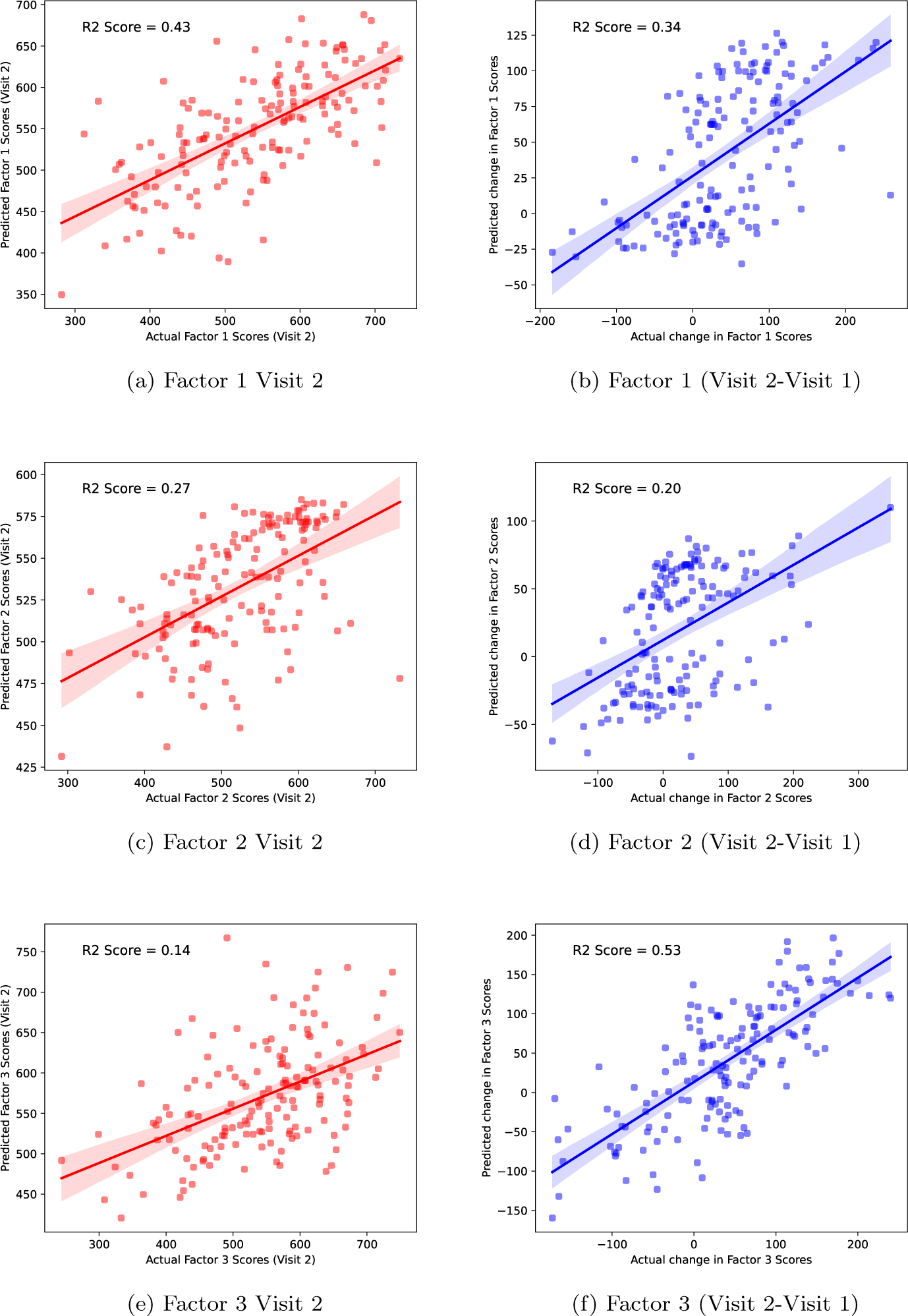
Regression plots illustrating predictions for factors 1, 2, and 3 using the image-assisted model framework. These plots are divided into two sets: those presented on the left side pertain to forecasting the factor scores specifically for Visit 2, whereas the plots on the right side focus on estimating the change in factors between Visit 2 and Visit 1.

### 3.2. Boxplot Analysis Emphasizing Statistical Significance of the Model

The boxplot in Fig. 4 displays the distribution of differences between predicted and test values of factors 1, 2, and 3 for the proposed two cases. Each box represents the interquartile range (IQR) of the differences, with the median marked by a horizontal line within the box. The whiskers extend to the minimum and maximum values within 1.5 times the IQR from the first and third quartiles. Any points outside this range are considered outliers and marked as individual points. Here the dashed red line indicates the mean difference value. This plot is useful for understanding the spread and central tendency of the prediction errors. If the boxplot is centered around zero with small variability, it suggests the model’s predictions are generally accurate. On the other hand, if the boxplot is skewed or has large variability, it indicates systematic errors or inconsistent predictions. Fig.4 (a), (c), and (e) illustrate differences between predicted and actual values for factor scores 1, 2, and 3 at Visit 2. We can observe that all three boxplots are centered around zero. Factor 1 exhibits the lowest variability, followed by the other two factors. Conversely, in Fig.4 (b), (d), and (f), Factor 3 demonstrates the least variability, followed by Factors 1 and 2. Across all 6 figures, the mean difference falls within the narrow range of -0.75 to 0.75, indicating that predicted values closely match actual values. This demonstrates the model’s accuracy with a consistent level of error across all cases.

**Figure 4:**
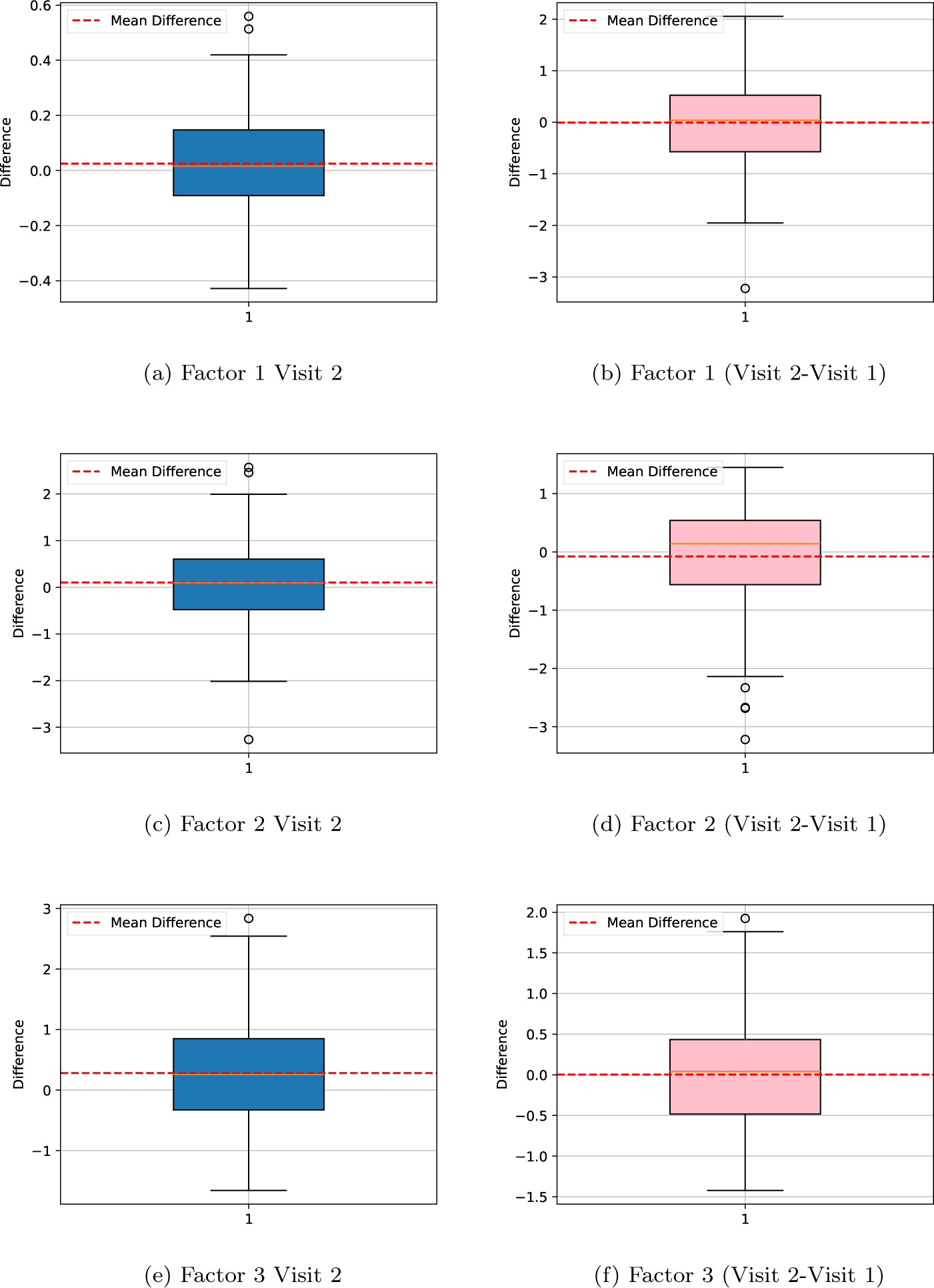
Boxplots illustrating the distribution of differences between predicted and actual values for factors 1, 2, and 3 using the image-assisted model framework. The plots are split into two groups: those on the left display differences between predicted and actual values for factor scores at Visit 2, while those on the right show changes in factors between Visit 2 and Visit 1.

Table. 4 uses statistical measures to compare the predicted values with the actual factor values, particularly focusing on the differences between them. The mean difference represents the average difference between the predicted and actual values. A positive mean difference indicates that the predictions tend to be higher than the actual values, while a negative mean difference indicates the opposite. A mean difference close to zero indicates that, on average, the predicted and actual labels are similar. Meanwhile, the standard deviation of differences measures the variability of the differences between predicted and actual labels. A higher standard deviation indicates greater variability in the differences. Notably, the PCVAE model consistently exhibits smaller mean differences and standard deviations across all cases. Regarding p-values, a large p-value (typically > 0.05) indicates that the observed differences between predicted and actual labels are not statistically significant, implying a close match between the model’s predictions and the actual values. For all factors, the PCVAE model yields low t-values and high p-values (above 0.05), indicating good performance in predicting the outcomes.

**Table 4:** Statistical evaluation of the PCVAE model using various metrics.

| Factors | Prediction | Mean difference<br>between actual and<br>predicted factors | Standard deviation<br>of differences between<br>actual and predicted factors | t-value | p-value |
| --- | --- | --- | --- | --- | --- |
| Factor 1 | Factor 1 Visit 2 | 0.02 | 0.16 | 1.87 | 0.06 |
|  | Factor 1<br>(Visit 2 - Visit 1) | -0.006 | 0.81 | -0.10 | 0.91 |
| Factor 2 | Factor 2 Visit 2 | 0.10 | 0.85 | 1.49 | 0.13 |
|  | Factor 2<br>(Visit 2 - Visit 1) | -0.07 | 0.89 | -1.08 | 0.28 |
| Factor 3 | Factor 3 Visit 2 | 0.28 | 0.88 | 1.98 | 0.32 |
|  | Factor 3<br>(Visit 2 - Visit 1) | 0.002 | 0.68 | 0.04 | 0.96 |

### 3.3. Comparitive Analysis of Model Performance Using R2 Scores

While considering the R2 scores across the assessment-only model, image-assisted model, and neuroimaging-only model for Visit 2 factor prediction, it becomes evident that the image-assisted PCVAE model consistently out-performs the rest of the models. This indicates that incorporating image data into the predictive model significantly enhances its predictive capabilities compared to relying solely on assessment data or neuroimaging data. The image-assisted PCVAE model achieves the highest R2 scores for all factors, suggesting that the additional information extracted from images plays a crucial role in improving prediction accuracy. These R2 score comparisons are illustrated in Fig. 5 for a clearer visual representation of the model performance differences. For factor 1, the R2 score increases from 0.38 in the SVR model to 0.42 in the PCVAE model, signifying a notable improvement in predictive accuracy. Similarly, for factor 2, the R2 score jumps from 0.10 to 0.27, demonstrating a substantial enhancement in predictive capability by employing the image-assisted model. Although factor 3 shows a slight increase in R2 score from 0.11 to 0.14, the improvement is less pronounced compared to factors 1 and 2. On the other hand, the neuroimaging-only model consistently performs the worst among the three models. The poor performance of the neuroimaging-only model could be attributed to several factors, one of which might be the low correlation between the sFNC data from Visit 1 and the predicted factors at Visit 2. This could be due to the limited scope of information provided by neuroimaging data alone, which may not capture the complexity and details of factors being predicted as effectively as the combined information from assessments and images.

**Figure 5:**
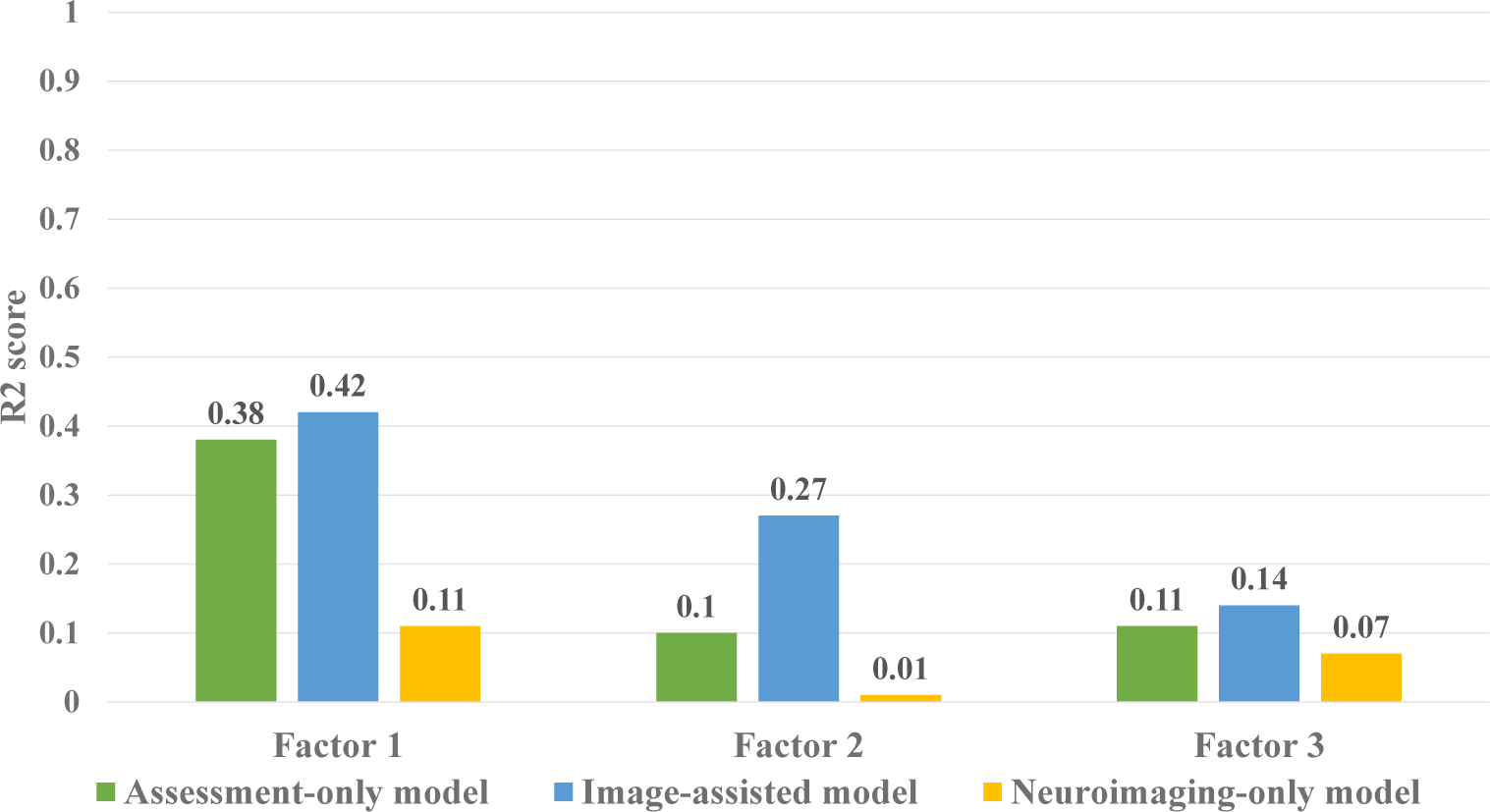
The variation in R2 scores across factors 1, 2, and 3 between the assessment-only model, image-assisted model, and neuroimaging-only model for Visit 2 factor prediction. Higher R2 values are indicative of better performance. These findings imply that the image-assisted model has assimilated latent information from neuroimaging data during training, potentially enhancing its predictive capabilities when solely relying on assessment data in future predictions.

Moreover, the significant performance gap observed, particularly in Factor 2, between the models underscores the importance of incorporating diverse data modalities for accurate prediction. In this case, the image-assisted PC-VAE model, by leveraging both assessment and image data, likely captures a broader range of features and patterns associated with Factor 2, leading to higher predictive accuracy.

Similarly, when examining the R2 scores across the three models for predicting changes in factors, the image-assisted PCVAE model surpasses the others across all factors. Notably, the neuroimaging-only model demonstrates the second-best performance for Factor 1 and Factor 2. For Factor 1, the R2 score increases from 0.27 in the RF model to 0.34 in the PCVAE model. While the image-assisted PCVAE model excels in predicting all three factors, the neuroimaging-only model exhibits competitive performance, particularly for Factors 1 and 2. This implies that the change in sFNC data carries substantial predictive significance, even in the absence of supplementary in-formation from assessment data. For Factor 2, the R2 score increases from 0.01 to 0.20 when comparing the SVR and PCVAE models, while the RF model achieves an R2 score of 0.15. Conversely, Factor 3 exhibits a modest increase in the R2 score from 0.50 to 0.53 when comparing the SVR and PCVAE models, although this improvement is less significant compared to Factors 1 and 2. These comparisons are visualized in Fig. 6, providing a bar graph representation of the performance differences among the models for each factor.

**Figure 6:**
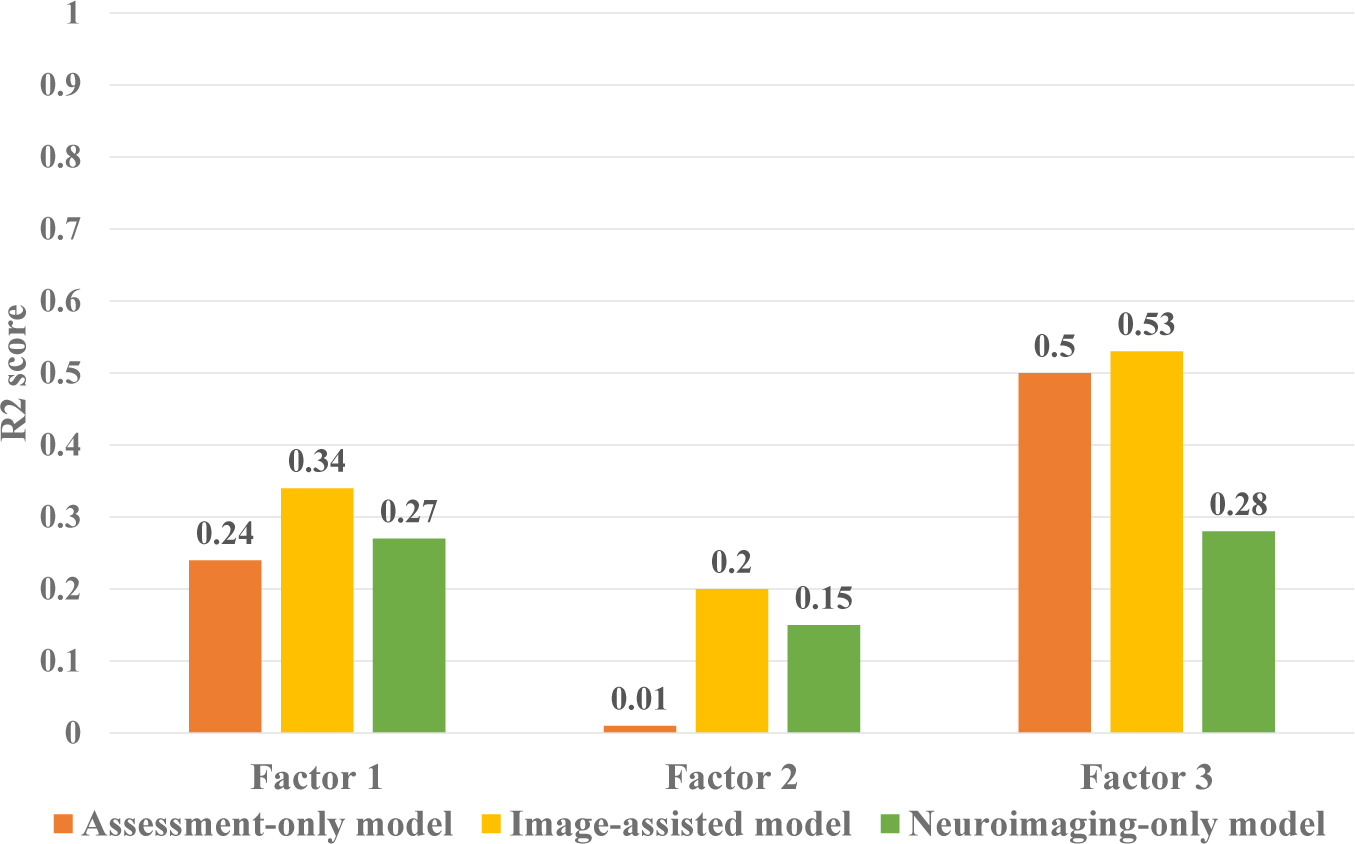
The variation in R2 scores across factors 1, 2, and 3 between the assessment-only model, image-assisted model, and neuroimaging-only model for change in factor (Visit 2 -Visit 1) prediction.

## 4. Discussion

The study aims to identify connections between neuroimaging data and corresponding assessments while concurrently predicting future behavior. Additionally, this work also captures insights about dimensions of brain health from neuroimaging data that may not be accessible through assessment data alone. This parallel exploration seeks to enhance our understanding of the complex relationship between neural patterns and behavioral measurements, offering a more comprehensive approach to predicting future behaviors. The neuroimaging analysis conducted in this study yielded crucial insights into the associated brain connectivity patterns. Essentially, the connectogram analysis played a pivotal role in visually showcasing these intricate network relationships. During the connectogram analysis of the connectedness factor, we observed positive correlations between cognitive control-subcortical domains, indicating enhanced connectivity within brain regions involved in decision-making and cognitive processing. Conversely, negative correlations were found in the cognitive control-sensorimotor and cognitive control-visual domains. The contributors to the connectedness factor, as identified by the factor-based model, include happiness, resilience, satisfaction, meaningful activities, social support, compassion, and social engagement. The characterization of the connectedness factor reveals the significant influence of positive psychological states and social factors on brain health, as indicated by the observed correlations and identified contributors.

Meanwhile, the emotional balance factor reflects the combination of positive (happiness) and negative (depression, anxiety, and stress) emotional states. The overlap in positive correlations with the connectedness factor suggests shared neural mechanisms between the two factors, particularly in cognitive control-subcortical domains. Moreover, the connectogram analysis of this factor also shows enhanced connectivity between visual processing and sensorimotor functions, indicating improved coordination between visual perception and motor functions. On the contrary, strong negative correlations were observed between the cognitive control-visual domains. Consequently, this factor represents a balanced emotional state with an emphasis on cognitive flexibility and emotional regulation. Finally, the clarity factor encompasses elements like strategic attention, innovation, fluidity of abstract interpretations (or possibility thinking), memory, sleep, outlook, and compassion. The corresponding connectogram showed a positive correlation between the auditory-default mode domain and the sensorimotor-visual domains. This indicates enriched cognitive and sensory-motor processing features of this factor.

In conclusion, while the assessment measures provide valuable information about components of brain health (i.e., cognition, emotional well-being, and social connectedness), the connectogram inferences reveal common functionalities. They also uncover hidden information that complements and extends the insights gained from assessment measures alone. The integration of both types of data provides a more comprehensive view of the complex relationships between behavioral performance across brain health domains and neural connectivity. Moreover, the post-intervention prediction performance is a crucial aspect of the study, emphasizing the practical implications of the neuroimaging findings. The substantial improvement in predictive accuracy with the PCVAE model, especially when incorporating sFNC data, underscores the significance of integrating advanced modeling techniques with neuroimaging information. The comparison between assessment-only (SVR), neuroimaging-only (RF) and image-assisted (PCVAE) models further highlights the importance of leveraging a broader range of information sources. The consistently superior performance of the image-assisted model across all factors indicates that multimodal data significantly improves predictive accuracy. The statistical evaluation also compared predicted and actual factor values, revealing that the PCVAE model consistently exhibits smaller mean differences and standard deviations. Additionally, PCVAE demonstrates low t-values and high p-values (> 0.05) across all factors, indicating effective prediction performance. Furthermore, the tighter clustering of regression lines and higher R2 scores in the PCVAE-based model suggests a more robust and consistent relationship between actual and predicted scores, enhancing the overall reliability of the predictive modeling approach.

## 5. Conclusion and Future Works

The research presents a novel predictive framework that integrates assessment data with neuroimaging insights, providing a holistic understanding of the relationship between neural patterns and behavioral outcomes. The inclusion of sFNC data in the model plays a pivotal role in advancing our understanding of how to validate the multidimensional aspects of brain health. Here the importance lies in the fact that brain health is a complex interconnection of various factors, and traditional assessment methods may not fully capture the subtleties of neural dynamics. The sFNC data provides a more detailed view of how different brain regions communicate and coordinate, offering insights into the underlying neural mechanisms associated with changes in brain health over time, either improvement or degradation. The enhanced predictive accuracy of this model opens avenues for identifying subtle changes in neural connectivity patterns that may precede observable behavioral outcomes. This, in turn, has potential implications for early detection of positive or negative changes in brain health. Moreover, these findings motivate combined data approaches to evaluate intervention effectiveness to promote brain health. In short, further research that explores combining longitudinal behavioral with neural data may offer a deeper understanding of the early neural basis of changes in one’s brain health trajectory at an individual level, perhaps making it possible to intervene proactively at earlier time points.

However, addressing limitations such as sample size constraints and reliance on self-reported assessment data will be essential for advancing the generalizability and robustness of this model. Future research should include longitudinal exploration, external validation across diverse populations, and the incorporation of additional modalities to further refine the understanding of how neural connectivity correlates with behavioral changes over time. By overcoming these challenges and building upon the current insights, this study lays the foundation for a comprehensive predictive framework that holds the potential for guiding interventions and promoting mental well-being.

## Abbreviations

AUD: Auditory
BHI: Brain Health Index
CB: Cerebellar
CC: Cognitive-Control
DM: Default Mode
GSP: Genomics Superstructure Project
HCP: Human Connectome Project
ICA: Independent Component Analysis
IC: Independent Component
Leaky ReLU: Leaky Rectified Linear Unit
MAE: Mean Absolute Error
MSE: Mean Square Error
MMSE: Mini-Mental Status Examination
PCVAE: Partially Conditional Variational Autoencoder
R2: Coefficient of Determination
RBF: Radial Basis Function
RF: Random Forest
rs-fMRI: Resting-State Functional Magnetic Resonance Imaging
RSN: Resting-State Networks
SM: Sensory-Motor
sFNC: Static Functional Network Connectivity
SC: Subcortical
SVR: Support Vector Regression
TMT: Trail Making Test
VIS: Visual
WHO: World Health Organization

## Credit authorship contribution statement

**Meenu Ajith** Conceptualization, Methodology, Software, Manuscript Writing & Editing. **Jeffrey S. Spence** Supervision, Funding Acquisition, Manuscript Writing & Editing. **Sandra B. Chapman** Supervision, Fund-ing acquisition, Manuscript Writing & Editing. **Vince D. Calhoun** Su-pervision, Conceptualization, Funding Acquisition, Manuscript Writing & Editing.

## Acknowledgements

This work was supported in part by the UTD Sammon’s award and NIH R01MH123610.

## References

Allen, T.T., Ashmore, L., Gordon, S., Tate, A., Cook, L.G., Chapman, S.B., 2020. Charisma™: A virtual reality training to promote social brainhealth in adults, in: Social Skills Across the Life Span. Elsevier, pp. 295–309.

Arevalo-Rodriguez, I., Smailagic, N., i Figuls, M.R., Ciapponi, A., Sanchez-Perez, E., Giannakou, A., Pedraza, O.L., Cosp, X.B., Cullum, S., 2015. Mini-mental state examination (mmse) for the detection of alzheimer’s disease and other dementias in people with mild cognitive impairment (mci). Cochrane database of systematic reviews.

Avan, A., Hachinski, V., Learn, B.H., Group, A., Aamodt, A.H., Alessi, C., Ali, S., Alladi, S., Andersen, R., Anderson, K.K., Avan, A., et al., 2022. Brain health: Key to health, productivity, and well-being. Alzheimer’s & Dementia 18, 1396–1407.

Battista, P., Salvatore, C., Castiglioni, I., et al., 2017. Optimizing neuropsychological assessments for cognitive, behavioral, and functional impairment classification: a machine learning study. Behavioural neurology 2017.

Burckhardt, C.S., Anderson, K.L., Archenholtz, B., Hägg, O., 2003. The flanagan quality of life scale: Evidence of construct validity. Health and quality of life outcomes 1, 1–7.

Buysse, D.J., Reynolds III, C.F., Monk, T.H., Berman, S.R., Kupfer, D.J., 1989. The pittsburgh sleep quality index: a new instrument for psychiatric practice and research. Psychiatry research 28, 193–213.

Chapman, S., Robertson, I., Zientz, J., Eyre, H., Ling, G., D’Esposito, M., 2022. The neuroscience of brain health. Lifestyle medicine.

Chapman, S.B., Fratantoni, J.M., Robertson, I.H., D’Esposito, M., Ling, G.S., Zientz, J., Vernon, S., Venza, E., Cook, L.G., Tate, A., et al., 2021. A novel brainhealth index prototype improved by telehealth-delivered training during covid-19. Frontiers in Public Health, 182.

Du, Y., Fu, Z., Sui, J., Gao, S., Xing, Y., Lin, D., Salman, M., Abrol, A., Rahaman, M.A., Chen, J., et al., 2020. Neuromark: An automated and adaptive ica based pipeline to identify reproducible fmri markers of brain disorders. NeuroImage: Clinical 28, 102375.

Eakman, A.M., 2011. Convergent validity of the engagement in meaningful activities survey in a college sample. OTJR: Occupational Therapy Journal of Research 31, 23–32.

Guo, Y., 2022. A selective review of the ability for variants of the trail making test to assess cognitive impairment. Applied Neuropsychology: Adult 29, 1634–1645.

Hachinski, V., Avan, A., Gilliland, J., Oveisgharan, S., 2021. A new definition of brain health. The Lancet Neurology 20, 335–336.

Hanten, G., Li, X., Chapman, S.B., Swank, P., Gamino, J., Roberson, G., Levin, H.S., 2007. Development of verbal selective learning. Developmental Neuropsychology 32, 585–596.

Hills, P., 2002. Argyle, m. The Oxford happiness questionnaire: A compact scale for the measurement of psychological well-being. Personality and Individual Differences 33, 1073–1082.

Hu, H.Y., Ou, Y.N., Shen, X.N., Qu, Y., Ma, Y.H., Wang, Z.T., Dong, Q., Tan, L., Yu, J.T., 2021. White matter hyperintensities and risks of cognitive impairment and dementia: a systematic review and meta-analysis of 36 prospective studies. Neuroscience & Biobehavioral Reviews 120, 16– 27.

Johnson, L.K., 2018. The light triad scale: developing and validating a preliminary measure of prosocial orientation. Ph.D. thesis. The University of Western Ontario (Canada).

Jurca, R., Jackson, A.S., LaMonte, M.J., Morrow Jr, J.R., Blair, S.N., Wareham, N.J., Haskell, W.L., van Mechelen, W., Church, T.S., Jakicic, J.M., et al., 2005. Assessing cardiorespiratory fitness without performing exercise testing. American journal of preventive medicine 29, 185–193.

Kingma, D.P., Ba, J., 2014. Adam: A method for stochastic optimization. arXiv preprint arXiv:1412.6980.

Lee, A., Shah, S., Atha, K., Indoe, P., Mahmoud, N., Niblett, G., Pradhan, V., Roberts, N., Malouf, R.S., Topiwala, A., 2024. Brain health measurement: a scoping review. BMJ open 14, e080334.

Lira, B., O’Brien, J.M., Peña, P.A., Galla, B.M., D’Mello, S., Yeager, D.S., Defnet, A., Kautz, T., Munkacsy, K., Duckworth, A.L., 2022. Large studies reveal how reference bias limits policy applications of self-report measures. Scientific Reports 12, 19189.

Lovibond, P.F., Lovibond, S.H., 1995. The structure of negative emotional states: comparison of the depression anxiety stress scales (dass) with the beck depression and anxiety inventories. Behaviour research and therapy 33 3, 335–43.

Maas, A.L., Hannun, A.Y., Ng, A.Y., et al., 2013. Rectifier nonlinearities improve neural network acoustic models, in: Proc. icml, Atlanta, GA. p. 3.

Nemesure, M.D., Heinz, M.V., Huang, R., Jacobson, N.C., 2021. Predictive modeling of depression and anxiety using electronic health records and a novel machine learning approach with artificial intelligence. Scientific reports 11, 1980.

Organization, W.H., et al., 2022. Optimizing brain health across the life course: Who position paper.

Schwarzer, R., Jerusalem, M., 1995. Generalized self-efficacy scale. J. Weinman, S. Wright, & M. Johnston, Measures in health psychology: A user’s portfolio. Causal and control beliefs 35, 37.

Sherbourne, C.D., Stewart, A.L., 1991. The mos social support survey. Social science & medicine 32, 705–714.

Strauss, C., Taylor, B.L., Gu, J., Kuyken, W., Baer, R., Jones, F., Cavanagh, K., 2016. What is compassion and how can we measure it? a review of definitions and measures. Clinical psychology review 47, 15–27.

Vas, A.K., Chapman, S.B., Cook, L.G., 2015. Language impairments in traumatic brain injury: a window into complex cognitive performance. Handbook of clinical neurology 128, 497–510.

